# BertST: BERT-based Spatial Domain Identification in Patient Data

**DOI:** 10.64898/2026.07.04.736527

**Authors:** Gospel Ozioma Nnadi

## Abstract

Spatial transcriptomics enables the study of gene expression within its native tissue context, providing critical insights into cellular organization and microenvironment-driven biological processes. A key challenge in this field is spatial domain identification, which aims to partition tissue into coherent regions by jointly leveraging gene expression and spatial information. Existing approaches are predominantly based on Graph Neural Networks (GNNs), and approach based on Transformers particularly, Bidirectional Encoder Reppresentation Transformer (BERT) model for modelling both local and long-range dependencies remains largely unexplored.

In this work, we propose BERT for Spatial Transcriptomics (BertST), a transformer-based framework that reformulates spatial transcriptomics as a graph-to-text representation learning problem. Building upon the BERTwalk paradigm, we construct a task-specific multi-graph representation integrating spatial adjacency, pruned gene-expression similarity, and a fully connected gene-expression graph. This design enables the modelling of both local spatial structure and global molecular relationships. Random walks over these graphs are treated as sequences, allowing a BERT model to learn contextualised node embeddings.

To further enhance representation quality, we introduce a hierarchical multi-graph propagation strategy, where embedding refinement is performed sequentially: first on the fully connected graph to capture global structure, followed by the pruned graph to refine molecular relationships, and finally on the spatial graph to enforce local smoothness. This ordering ensures that global information is effectively distributed and progressively constrained by biologically meaningful neighbourhoods.

We also improve computational efficiency by leveraging *PecanPy*, a fast and scalable implementation of node2vec, enabling efficient random walk generation on dense graphs. Experimental results on multiple 10x Visium datasets, including DLPFC and Human Breast Cancer, demonstrate that BertST consistently outperforms or matches GNN-based methods such as ConST, CCST, and SpaceFlow in terms of Adjusted Rand Index (ARI) and Adjusted Mutual Information (AMI).

Overall, BertST highlights the potential of transformer-based architectures for spatial omics analysis by effectively capturing both local and long-range spatial-molecular dependencies, offering a promising alternative to traditional graph-based methods.

## 1 Introduction

Spatial transcriptomics has revolutionized the study of complex biological systems by enabling the measurement of gene expression while preserving the spatial organization of cells within tissues. In contrast to conventional transcriptomic approaches that treat tissues as homogeneous mixtures, spatial technologies provide a structured view of cellular organization, allowing researchers to investigate how cellular function is shaped by local microenvironments. This spatially resolved perspective has driven major advances in disease biology, including tumor heterogeneity, immune–tumor interactions, and disease progression, as well as in developmental biology through improved understanding of spatiotemporal gene regulation, morphogenesis, intercellular communication, and the emergence of distinct cell subpopulations [1].

A central computational challenge in this domain is the identification of spatially coherent cell subpopulations, commonly referred to as *spatial domain identification*. This task seeks to partition tissue into biologically meaningful regions by jointly leveraging gene expression profiles and spatial context. In recent years, this problem has attracted significant attention, with deep learning-based methods—particularly Graph Neural Networks (GNNs)—emerging as the dominant paradigm due to their ability to model spatial relationships between cells through message passing.

Representative methods include ConST [2], which integrates gene expression, spatial coordinates, and histological morphology through contrastive learning. It employs masked autoencoders (MAE) for morphological feature extraction, applies Principal Component Analysis (PCA) for dimensionality reduction, and adopts a two-stage training strategy combining reconstruction and representation alignment. CCST [3] incorporates spatial and molecular information into unified graph representations and leverages Deep Graph Infomax (DGI) [4] to learn latent embeddings. Similarly, SpaceFlow [5] utilizes graph neural networks with spatial regularization to generate compact embeddings that preserve both transcriptomic and spatial structure.

Despite their success, these GNN-based approaches are inherently local message-passing mechanisms. In contrast, transformer-based architectures remain largely underexplored in spatial transcriptomics to capture local and long-range dependencies and higher-order contextual relationships across tissue regions. Models such as Bidirectional Encoder Representations from Transformers (BERT), originally developed for natural language processing, learn rich contextual representations through self-supervised masked language modeling over large corpora [6]. Their capacity to model global dependencies makes them particularly well-suited for spatial biology, where cellular behaviour is governed by both local interactions and distal regulatory effects.

To address this gap, we propose BERT for Spatial Transcriptomics (BertST) (Figure 1), a transformer-based framework for spatial domain identification. Building upon the graph-to-text paradigm introduced in BERTwalk [7], we reformulate spatial transcriptomics as a sequence modeling problem. Specifically, we construct a multi-graph representation in which nodes correspond to spatial spots and edges encode spatial proximity and gene expression similarity. Random walks over these graphs generate sequences that serve as a corpus for BERT pretraining, enabling the model to learn joint spatial–molecular representations.

**Figure 1:**
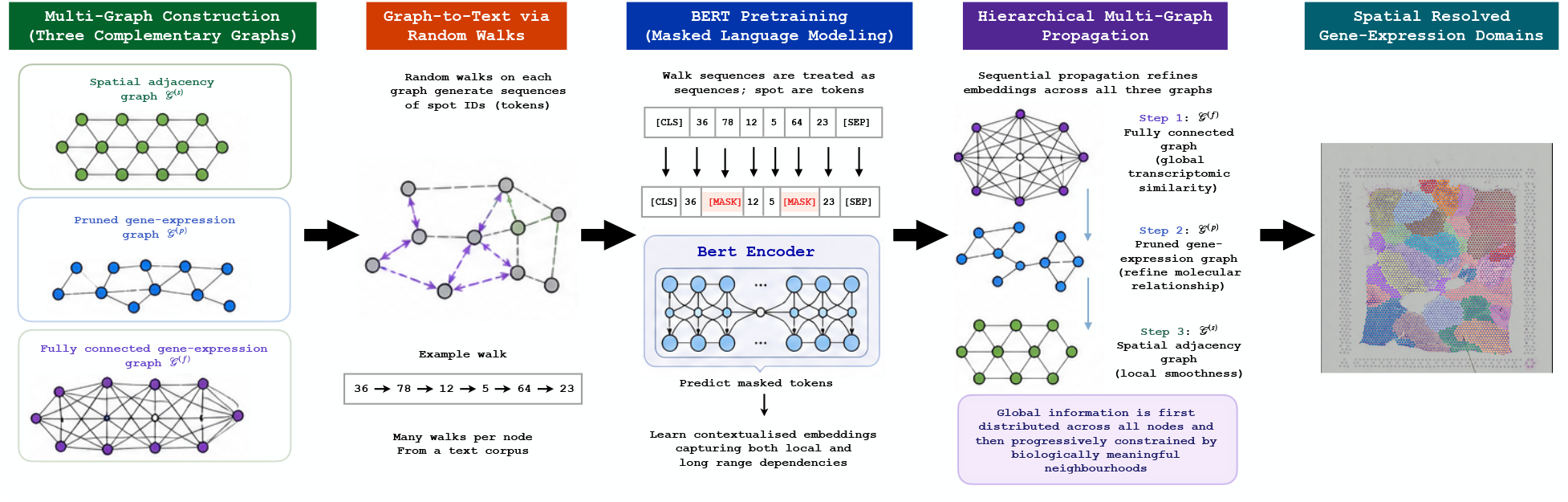
Overview of BertST for spatial domain identification. The method transforms graph-structured spatial transcriptomics data into sequences via random walks and leverages a BERT-based model to learn contextual representations, enabling accurate identification of spatial domains.

A key contribution of BertST lies in its task-specific multi-graph formulation, which integrates three complementary graph structures: a spatial adjacency graph capturing local tissue architecture, a pruned gene-expression graph encoding high-confidence molecular similarity, and a fully connected gene-expression graph that enables global interactions across all nodes. Crucially, the fully connected graph allows information to propagate across the entire tissue, enabling the modelling of long-range dependencies.

Beyond graph construction, BertST introduces a hierarchical propagation strategy that further refines learned representations. In particular, propagation is performed sequentially across the three graphs: first on the fully connected graph to capture global structure, followed by the pruned graph to refine molecular relationships, and finally on the spatial graph to enforce local smoothness. This ordering is critical, as it ensures that global information is first distributed across all nodes and then progressively constrained by biologically meaningful neighbourhoods, resulting in representations that are both globally consistent and locally coherent.

In addition, we improve the efficiency and scalability of graph-based sequence generation by leveraging *PecanPy* [8], a fast and parallelised implementation of node2vec. This enables efficient random walk sampling over dense graphs, which is essential for incorporating global similarity structure in spatial transcriptomics data.

We evaluate BertST on multiple spatial transcriptomics datasets generated using the 10x Visium platform [9]. Experimental results demonstrate that BertST consistently outperforms strong graph-based baselines, including ConST, CCST, and SpaceFlow, across standard clustering metrics such as Adjusted Rand Index (ARI) [10] and Adjusted Mutual Information (AMI) [11]. These findings highlight the effectiveness of transformer-based architectures for modeling complex spatial and molecular dependencies and suggest that language-model-inspired approaches provide a promising new direction for spatial omics analysis.

## 2 Methods

BertST (Figure 1) is a transformer-based framework for spatial domain identification that integrates spatial proximity and gene expression similarity through a graph-to-text representation learning paradigm. The method builds upon the BERTwalk strategy [7], adapting it to spatial transcriptomics by introducing a tailored graph formulation and efficient random walk generation. In particular, we leverage *PecanPy* [8] for scalable and parallelised node2vec-style sampling, and design a multi-graph representation that enables both local and global information propagation.

### 2.1 Data Preprocessing

Gene expression counts are library-size normalised, log-transformed, and restricted to the top 3,000 highly variable genes. Principal Component Analysis (PCA) is then applied to obtain a compact representation **Z**^(0)^∈ℝ^*N ×d*^ with *d* = 20. Spots without spatial neighbours are removed to ensure graph connectivity.

### 2.2 Multi-graph Graph Construction

A central contribution of BertST lies in its task-specific multi-graph formulation, which integrates spatial structure and gene expression similarity into a unified representation. Let *G*= *{G*^(*f*)^, *G*^(*p*)^, *G*^(*s*)^*}* denote the set of three graphs constructed over the same set of nodes *V* = *{v*_1_, …, *v*_*N*_ *}*, where each node corresponds to a spatial spot.

#### Gene expression similarity graph

Given PCA-reduced features **Z** ∈ R^*N ×d*^, we compute pairwise cosine similarity:

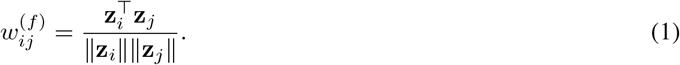

To ensure non-negativity, similarities are shifted:

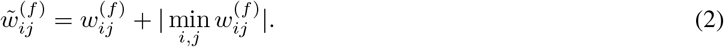

This defines the **fully connected graph *G***^(*f*)^ = (*V, ε*^(*f*)^), where all node pairs with 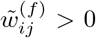 are connected. This graph encodes global molecular similarity and enables long-range interactions across all spots.

### Pruned gene expression graph

To retain only the most informative molecular relationships, we construct a *k*-nearest neighbour graph:

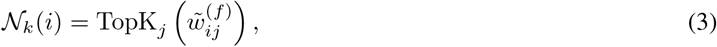

and define edges only for *j* ∈ *N*_*k*_(*i*) with *k* = 6. This yields the pruned graph^(*p*)^, which captures high-confidence gene similarity while reducing noise.

### Spatial graph with interaction refinement

Spatial adjacency is first defined using the Visium hexagonal grid:

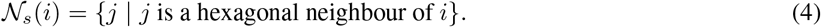

We further refine this graph by incorporating both Euclidean proximity and gene similarity. A *k*-nearest spatial graph is constructed:

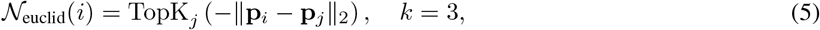

where **p**_*i*_ denotes spatial coordinates.

Additionally, a best-neighbour clustering strategy links spatial neighbours with minimal cosine distance:

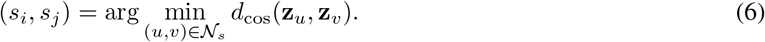

The final spatial adjacency matrix is defined as:

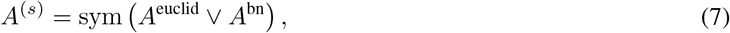

where *V* denotes element-wise union and sym(*·*) enforces symmetry.

The three graphs serve complementary roles:

- *G*^(*f*)^: global structure and long-range dependencies,
- *G*^(*p*)^: high-confidence molecular neighbourhoods,
- *G*^(*s*)^: local spatial coherence.

This design ensures that BertST captures hierarchical relationships, from global similarity to local spatial organisation, allowing information learned during training to propagate beyond immediate neighbours, capturing long-range dependencies that are essential in spatial biology.

### 2.3 Graph-to-Text Transformation via Random Walks

Following the BERT walk strategy [7], we recast the multi-graph *G* as a text corpus by generating random walk sequences and treating each node as a token.

Unlike the original implementation, we leverage *PecanPy* [8], an efficient and parallelised node2vec framework, to perform scalable random walk sampling on dense graphs. Given a node *v*_*i*_, we generate *r* walks of length *l*:

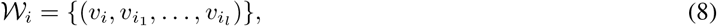

where transitions follow a second-order biased random walk.

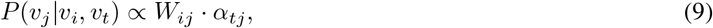

where *α*_*tj*_ controls return and exploration biases. The use of multiple graphs during walk generation enables sequences that encode both spatial adjacency and global similarity structure

### 2.4 BERT-Based Representation Learning

The resulting corpus *C*= U_*i*_ *W*_*i*_ is used to pretrain a BERT model with a masked language modelling objective [12], enabling the model to learn contextualised node representations that integrate spatial and molecular context.

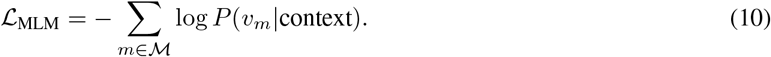

By training on sequences derived from multiple graphs, BertST captures both local and global dependencies.

### 2.5 Multi-Graph Propagation

To further enhance representation consistency, BertST applies a sequential propagation mechanism across the three graphs. Let **E**^(0)^ denote the initial BERT embeddings. For each graph *G*^(*k*)^, we perform propagation using APPNP:

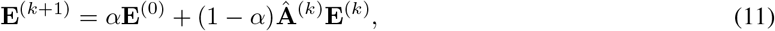

where Â^(*k*)^ is the normalised adjacency matrix of graph *G*^(*k*)^.

Importantly, propagation is applied sequentially over the three graphs:

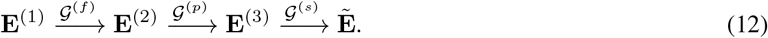

This ordering is critical: propagation begins with the fully connected graph to capture global structure, followed by the pruned graph to refine molecular relationships, and finally the spatial graph to enforce local smoothness. This hierarchical propagation ensures that global information is first distributed across all nodes and then progressively constrained by biologically meaningful neighbourhoods.

Overall, this multi-graph propagation mechanism allows BertST to effectively integrate global and local dependencies, overcoming limitations of purely local message-passing approaches.

### 2.6 Clustering

The final embeddings are clustered using mClust [13] to obtain spatial domains. Performance is evaluated using Adjusted Rand Index (ARI) and Adjusted Mutual Information (AMI) against ground truth annotations.

## 3 Results

We applied BertST on publicly available dataset such as the Dorsolateral Prefrontal Cortex (DLPFC) twelve datasets, and the Human breast cancer (HBC) dataset from Visium. BertST perform was evaluated using ARI and AMI.

### 3.1 Overall Performance on DLPFC

Table 1 reports the clustering performance across the 12 DLPFC datasets using ARI and AMI. Overall, BertST achieves the best average performance among all three all GNN-based baselines methods, with a mean ARI of 0.45 and a mean AMI of 0.60, outperforming.

**Table 1:**
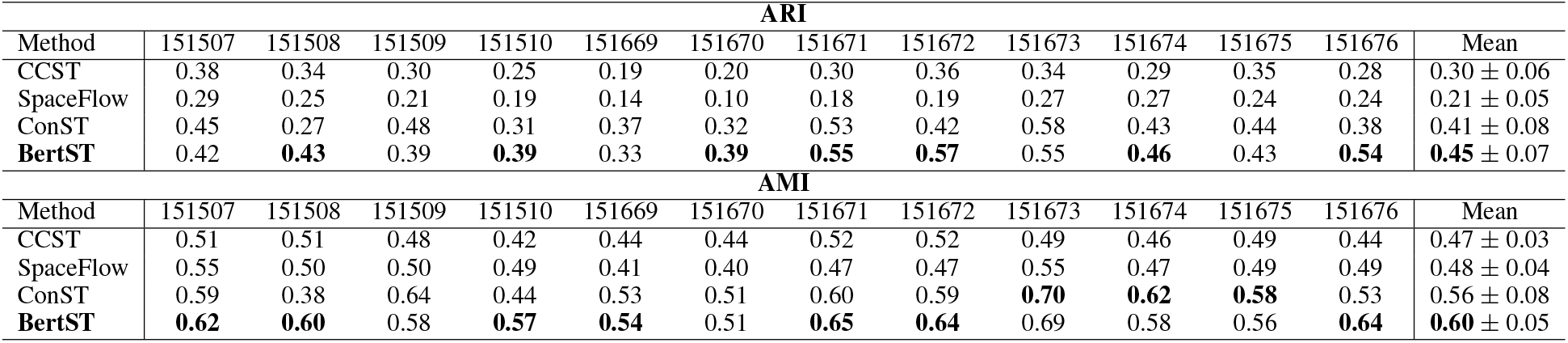
Clustering performance on DLPFC datasets using Adjusted Rand Index (ARI) and Adjusted Mutual Information (AMI). Best results are shown in bold.

| ARI |  |  |  |  |  |  |  |  |  |  |  |  |  |
| --- | --- | --- | --- | --- | --- | --- | --- | --- | --- | --- | --- | --- | --- |
| Method | 151507 | 151508 | 151509 | 151510 | 151669 | 151670 | 151671 | 151672 | 151673 | 151674 | 151675 | 151676 | Mean |
| CCST | 0.38 | 0.34 | 0.30 | 0.25 | 0.19 | 0.20 | 0.30 | 0.36 | 0.34 | 0.29 | 0.35 | 0.28 | $0.30 \pm 0.06$ |
| SpaceFlow | 0.29 | 0.25 | 0.21 | 0.19 | 0.14 | 0.10 | 0.18 | 0.19 | 0.27 | 0.27 | 0.24 | 0.24 | $0.21 \pm 0.05$ |
| ConST | 0.45 | 0.27 | 0.48 | 0.31 | 0.37 | 0.32 | 0.53 | 0.42 | 0.58 | 0.43 | 0.44 | 0.38 | $0.41 \pm 0.08$ |
| **BertST** | 0.42 | **0.43** | 0.39 | **0.39** | 0.33 | **0.39** | **0.55** | **0.57** | 0.55 | **0.46** | 0.43 | **0.54** | **0.45** $\pm 0.07$ |
| AMI |  |  |  |  |  |  |  |  |  |  |  |  |  |
| Method | 151507 | 151508 | 151509 | 151510 | 151669 | 151670 | 151671 | 151672 | 151673 | 151674 | 151675 | 151676 | Mean |
| CCST | 0.51 | 0.51 | 0.48 | 0.42 | 0.44 | 0.44 | 0.52 | 0.52 | 0.49 | 0.46 | 0.49 | 0.44 | $0.47 \pm 0.03$ |
| SpaceFlow | 0.55 | 0.50 | 0.50 | 0.49 | 0.41 | 0.40 | 0.47 | 0.47 | 0.55 | 0.47 | 0.49 | 0.49 | $0.48 \pm 0.04$ |
| ConST | 0.59 | 0.38 | 0.64 | 0.44 | 0.53 | 0.51 | 0.60 | 0.59 | **0.70** | **0.62** | **0.58** | 0.53 | $0.56 \pm 0.08$ |
| **BertST** | **0.62** | **0.60** | 0.58 | **0.57** | **0.54** | 0.51 | **0.65** | **0.64** | 0.69 | 0.58 | 0.56 | **0.64** | **0.60** $\pm 0.05$ |

In terms of ARI, BertST consistently ranks among the top-performing methods and achieves the highest scores in the majority of datasets, including 151508, 151510, 151670, 151671, 151672, 151674, and 151676. Notably, BertST shows substantial improvements over ConST in several sections, particularly in 151508 and 151672, indicating stronger alignment with ground truth spatial domains. While ConST performs competitively on specific datasets such as 151673, BertST demonstrates more stable performance across all samples, leading to a higher overall mean.

A similar trend is observed for AMI, where BertST achieves the highest average score and outperforms competing methods in most datasets, including 151507, 151508, 151510, 151669, 151671, 151672, and 151676. Although ConST attains the best AMI in a few cases (e.g., 151673–151675), BertST maintains consistently strong performance, suggesting improved robustness in capturing both cluster purity and completeness.

Compared to CCST and SpaceFlow, which exhibit lower mean performance and higher variability, BertST demonstrates clear advantages in both accuracy and stability. These results indicate that transformer-based contextual modeling enables more effective integration of spatial and transcriptomic information than conventional graph-based message passing alone.

### 3.2 Performance on Human Breast Cancer Dataset

Table 2 summarizes the results on the Human Breast Cancer (HBC) dataset. BertST achieves an ARI of 0.46, matching the best-performing method (CCST), and outperforming both SpaceFlow (0.42) and ConST (0.44). This indicates that BertST is able to accurately recover spatial domain structures in complex tumor tissue.

**Table 2:** Clustering performance on the Human Breast Cancer (HBC) dataset using Adjusted Rand Index (ARI) and Adjusted Mutual Information (AMI). Best results are shown in bold.

| Method | ARI | AMI |
| --- | --- | --- |
| CCST | **0.46** | **0.69** |
| SpaceFlow | 0.42 | 0.67 |
| ConST | 0.44 | 0.59 |
| BertST | **0.46** | 0.66 |

In terms of AMI, BertST obtains a score of 0.66, which is competitive with SpaceFlow (0.67) and slightly below CCST (0.69), while substantially outperforming ConST (0.59). Although BertST does not achieve the highest AMI on this dataset, its strong ARI performance suggests that it produces cluster assignments that are well aligned with ground truth labels.

### 3.3 Discussion

Across both DLPFC and HBC datasets, BertST demonstrates competitive and often superior performance compared to existing GNN-based approaches. The improvements are particularly pronounced in layered tissues such as the DLPFC benchmark, where BertST achieves the highest average scores and exhibits greater consistency across multiple tissue sections. On the HBC dataset, BertST remains highly competitive, achieving top ARI performance and strong AMI results, confirming its robustness in heterogeneous and complex tissue environments.

These results validate the effectiveness of the proposed graph-to-text transformer framework, which builds upon the BERTwalk paradigm [7] and extends it to spatial transcriptomics. A key contribution of this work lies in the tailored multi-graph formulation, where spatial, pruned gene-expression, and fully connected gene-expression graphs are jointly utilised. In particular, the inclusion of the fully connected graph enables global information propagation across all nodes, allowing BertST to capture long-range dependencies.

Furthermore, the hierarchical propagation strategy plays a critical role in the overall performance. Specifically, propagation begins with the fully connected graph to capture global structure, followed by the pruned graph to refine molecular relationships, and finally the spatial graph to enforce local smoothness. This ordering is essential, as it ensures that global information is first distributed across all nodes and then progressively constrained by biologically meaningful neighbourhoods. Such a design enables BertST to effectively balance global coherence and local spatial consistency, which is fundamental for accurate spatial domain identification.

Another important contribution is the adoption of *PecanPy* [8] for efficient and scalable random walk generation. Compared to the original node2vec implementation used in BERTwalk, this significantly improves computational efficiency and enables effective sampling over dense graphs, which is critical for spatial transcriptomics applications. In addition, we introduce task-specific adaptations, including graph construction strategies and parameter tuning, to better reflect the characteristics of spatial-molecular data.

From a computational perspective, the overall training time is strongly influenced by the size of the generated text corpus, which is determined by the random walk procedure. The complexity scales with the number of nodes, the number of walks per node, the walk length, and the number of sampling repetitions. As these factors increase, the corpus grows proportionally, making random walk generation the primary computational bottleneck, particularly for large-scale datasets therefore, making its optimization through *PecanPy* [8] particulary essential.

By leveraging the contextual learning capabilities of BERT on graph-derived sequences, BertST learns representations that integrate both spatial neighbourhood structure and molecular similarity. This leads to improved spatial domain identification, as reflected in clustering accuracy and robustness. While GPU acceleration can be applied during transformer training, the overall end-to-end speedup is limited and requires high GPU memory to load the multi-graph during propagation. Nevertheless, GPU usage can provides practical reductions in total training time.

Despite these advances, there remains room for further improvement. Incorporating explicit structural learning objectives, such as those proposed in GIST [14], could further enhance the quality of the learned graph representations by enforcing spatial coherence during training. Another promising direction is to replace the current random-walk-based text generation strategy with a more biologically informed sequence generation process. Rather than relying solely on transition probabilities derived from graph edge weights, future approaches could incorporate biological priors, to generate graph-derived node sequences that better reflect underlying biological processes. Such biologically guided graph-to-text representations may enable the transformer to learn richer spatial–molecular context and further improve domain identification.

Beyond algorithmic improvements, evaluating spatial domain identification methods using a broader range of complementary metrics is equally important. While ARI and AMI remain the most widely adopted measures, they primarily assess agreement with ground-truth annotations. Evaluation frameworks such as MultimetricST [15], which combine multiple spatial, and biological quality metrics, could provide a more comprehensive and robust assessment of model performance.

## Data and Code availability

The spatial transcriptomics datasets are available at https://zenodo.org/records/19153414 and https://figshare.com/articles/dataset/Visium_DLPFC_preprocessed/22004273. BertST is also available on github https://github.com/gospelnnadi/BertST.

